# Genetic structure of sugar kelp in the St. Lawrence Estuary and Gulf (Québec, Canada) Genetic structure of sugar kelp

**DOI:** 10.1101/2025.04.01.646547

**Authors:** Marie Treillefort, Sabrina Le Cam, Myriam Valero, Stéphane Mauger, Paolo Ruggieri, Flora Salvo, Isabelle Gendron-Lemieux, Tamara Provencher, Rénald Belley, France Dufresne

**Affiliations:** Département de biologie, Université du Québec à Rimouski; CNRS, Sorbonne Université, PUCCH, UACH, UMI 3614, Evolutionary Biology and Evolution of Algae, Station Biologique de Roscoff, Roscoff, France; CNRS, Sorbonne Université, UMR 7144, Dispersal Speciation and Evolution of Marine Species, Laboratory Adaptation et Diversité en Milieu Marin (AD2M), Station Biologique de Roscoff, Roscoff, France; CNRS, La Rochelle Université, UMR7266, Littoral Environnement et Sociétés, La Rochelle, France; Xelect Ltd, Horizon House, Abbey Walk, KY16 9LB, St Andrews, United Kingdom; Merinov. 96 montée de Sandy Beach, Gaspé, QC; École des pêches et de l’aquaculture du Québec, Cégep de la Gaspésie et des Îles, 167 Grande-Allée E, Grande-Rivière; Fisheries and Oceans Canada, Maurice Lamontagne Institute, Mont-Joli, Québec, Canada

**Keywords:** Population genetics, Microsatellite markers, *Saccharina latissima*, DartSeq, Genomic markers, St.Lawrence Estuary and Gulf

## Abstract

The sugar kelp, *Saccharina latissima* is cultivated at low scale in Quebec, Canada and current practice involve seeding meiospores or gametophyte stocks onto spools carrying twine and transferring these to a seaweed farm site. As the stocks can originate from locations spanning several hundreds of kilometers from the farm sites, such practice could involve genetic contamination and disrupt local adaptations. Assessing genetic structure can inform of the potential risks associated with this practice. Here we characterized the genetic diversity and structure of *S. latissima* from locations in the St. Lawrence Estuary and Gulf at both microsatellite loci (308 sporophytes at 22 loci in 16 sites) and genomic markers (228 sporophytes at 6578 single nucleotide polymorphisms in 13 sites). Several populations had low heterozygosity values and significant FIS values at microsatellite loci. No genetic structure was found among populations with microsatellite loci but strong genetic structuring was found with the genomic data. Population structure followed a geographic pattern and was congruent with major currents. Individuals from the wild population in the vicinity of the farm site were genetically distinct from the sporophytes on the growing lines that belong to a genetically distinct group. There was no significant genetic differentiation between wild individuals living in proximity of the farm site and another wild population of the same area. Hence aquaculture practices have not resulted in changes in the genetic composition of the wild population at large scale. Our results are important to guide future conservation efforts and for the seaweed farming industry.

## Introduction

The sugar kelp *Saccharina latissima* (Linnaeus) C.E. Lane, C. Mayes, Druehl & G.W. Saunders (Lane *et al*. 2006) is a widely distributed boreal-temperate species forming an important component of kelp forests. These forests participate in wave attenuation and carbon sequestration as well as providing nurseries, shelters, and feeding sites for many marine species (Dayton, 1985; Gaines and Roughgarden, 1987; Shil, et al. 2023). Kelp farming has long been carried out in coastal Asia and the practice is expanding in Europe, South America, and North America (Bjerregaard et al. 2016; Tamigneaux and Johnson 2016; Augyte et al. 2017; Breton et al. 2018; Campbell et al. 2019; Grebe et al. 2019; Hwang et al. 2019; Kim et al. 2019; Goecke et al. 2020). *S. latissima* is currently one of the main cultivated species in Europe and North America (Tamigneaux and Johnson 2016; Breton et al. 2018). Kelp species are usually harvested when mature in the wild and grown on lines and harvested for use in many industries (pharmaceutical and biomedical products, cosmetics, feed supplements, or biofuels) (Bartsch et al., 2008; Forbord et al., 2012; Jahan et al., 2017; Marinho et al., 2015). The increased popularity of this species for seaweed farming raises the question of developing best practices including farming without perturbing wild populations.

*S. latissima* has been cultivated experimentally and at low scale in the St. Lawrence Gulf since 2016 (Tamigneaux and Johnson 2016). This culture involves collecting mature sorus from wild individuals and seeding young sporophytes on farming lines until sufficient growth is reached to harvest. Seedlings were then transferred on lines at sites located several hundreds of kilometers away from their original location. Even though sporophytes are usually harvested before developing fertile meiospores, there might be a risk of genetic contamination if sporophytes are detached from the culture lines and disperse in nearby wild populations. The risk of genetic contamination depends on the extent of genetic differentiation between the population on the cultivation lines and the wild populations near the algaculture sites. Addressing the genetic diversity and genetic structure of this important founding species in the St. Lawrence Estuary and Gulf is an important step towards an efficient management.

Kelp has a dispersal capacity through sporophytes and meiospores, from a few meters to hundreds of kilometers (Dayton, 1985; Billot et al. 2003; Reed et al. 1992) depending on several factors such as habitat connectivity and marine currents (Alberto et al. 2010; Coleman et al. 2011; Durrant et al. 2018; Valero et al. 2011; Jaugeon et al. submitted). Oceanographic processes appeared to be more important drivers than geographic distance for finer spatial resolution in populations of *Laminaria digitata* (Fouqueau et al. 2024). Many studies have examined the genetic structure of the sugar kelp *S. latissima* using different types of molecular markers reflecting different time scales (Breton et al. 2018; Guzinski et al. 2016; Møller Nielsen et al. 2016; Neiva et al. 2018). Deep biogeographic history has been revealed with mitochondrial DNA (Neiva et al. 2018). Three phylogroups corresponding to (1) NE (British Columbia) Pacific, Greenland and Hudson Bay, (2) NE Atlantic, and (3) NW Atlantic were characterized (Neiva et al. 2018). Colonization from the Northeast Pacific into the Arctic and the Northeast Atlantic is thought to have occurred 5.5 million years ago (Luttikhuizen et al. 2018). Pleistocene glaciations have compressed NE Atlantic populations to the south and isolated NE and NW Atlantic populations (Neiva et al. 2018). At the scale of the eastern Atlantic region, northern (Norway, Sweden, Denmark) and southern (France) populations formed distinct clusters at 25 microsatellite loci, with an average FST of 0.36 (Guzinski et al., 2016). Strong genetic structure has been detected at genomic markers among sites from Scotland to Portugal (Guzinski et al. 2020). Within-population genetic diversity was lowest for the southern populations (Spain and Portugal) and the isolated island population on Helgoland, German Bight, and highest in Spitsbergen for both single nucleotide polymorphism (SNP) and expressed-sequence-tag (EST)-derived microsatellites (Guzinski et al. 2020). A survey of 21 sites along the Norwegian coastline using 9 microsatellite loci showed a relatively uniform genetic structure along 1000 kms (Ribeiro *et al*., 2022). By contrast, fine-scale genetic structure and low within-population genetic diversity has been found using microsatellite markers along the Maine region, USA (Breton *et al*., 2018).

Throughout the entire Northern Hemisphere, populations of *S. latissima* have undergone extensive changes in abundance and depth, including both expansions and declines (Moy and Christie 2012; Filbee-Dexter *et al*. 2016; Casado-Amezúa *et al*. 2019; Diehl et al. 2024). In the St. Lawrence, this decline appears to be related to two main factors, the warming of the water (Bernier et al. 2018) which no longer allows for optimal growth (Smale 2020), and an increased abundance of native (sea urchins) grazers and invasive exotic colonial bryozoans (e.g., *Membranipora membranace*, encrusting epiphytic bryozoa) (Tamigneaux and Johnson 2016; Bernier et al. 2018). Despite its importance as a foundation species and for the seaweed farming industry, baseline levels of genetic and genomic diversity have never been recorded in this region. There have been few genetic studies covering both the Estuary and the Gulf of St. Lawrence. A recent genomic study on the northern shrimp has revealed that individuals from the Estuary did not differ genetically from those from the northern Gulf (Bourret et al. 2024). Genetic differentiation was found (very low FST values) among larger sampling regions: Estuary and northern St. Lawrence Gulf, Scotian Shelf, Newfoundland and Labrador, and the Flemish Cap at neutral SNP (Bourret et al. 2024). Surface temperature and salinity were two important abiotic factors explaining genetic variation of shrimps in these regions (Bourret et al. 2024). Another genomic study on a species with high dispersal, the Greenland halibut, found an absence of population differentiation between Canada and west Greenland, and a weak genetic differentiation between the Gulf of St. Lawrence and the remainder of the Northwest Atlantic (Ferchaud et al. 2022). No genetic differentiation was found between fish from the Estuary and the Gulf of St. Lawrence (Ferchaud et al. 2022).

This study aimed to characterize genetic diversity and assess genetic structure of wild populations of *S. latissima* from St. Lawrence seaway using microsatellite and genomic markers. As this species does not disperse actively, we expect currents to drive much of the genetic structure at neutral SNP. We expect that sites located on the North Shore will be differentiated from other sites because of the presence of the Anticosti gyre (Figure 1). The Gaspé current outflows from the St. Lawrence River, which moves around the Gaspé Peninsula and along the southern shore of the Gulf of St. Lawrence (Savenkoff et al. 1996). Therefore, sites in the Estuary and the Gaspé coast may form another genetic cluster. Finally, sites from the southern Gaspé coast and the southern Gulf should be genetically distinct due to the downward movement of the Gaspé current. As individuals of *S. latissima* may dislodge from lines during their cultivation, there may be genetic contamination by individuals grown from the Baie-des-Chaleurs in the North Shore population. Assuming that sugar kelps from these two areas are genetically distinct, genetic contamination would result in sporophytes from areas near the culture sites to be more genetically related to the Baie-des-Chaleurs sites than to populations from their own region. Information on this key species can help create appropriate recommendations for managing aquaculture and preserving biodiversity in the St. Lawrence Gulf and Estuary.

**Figure 1.**
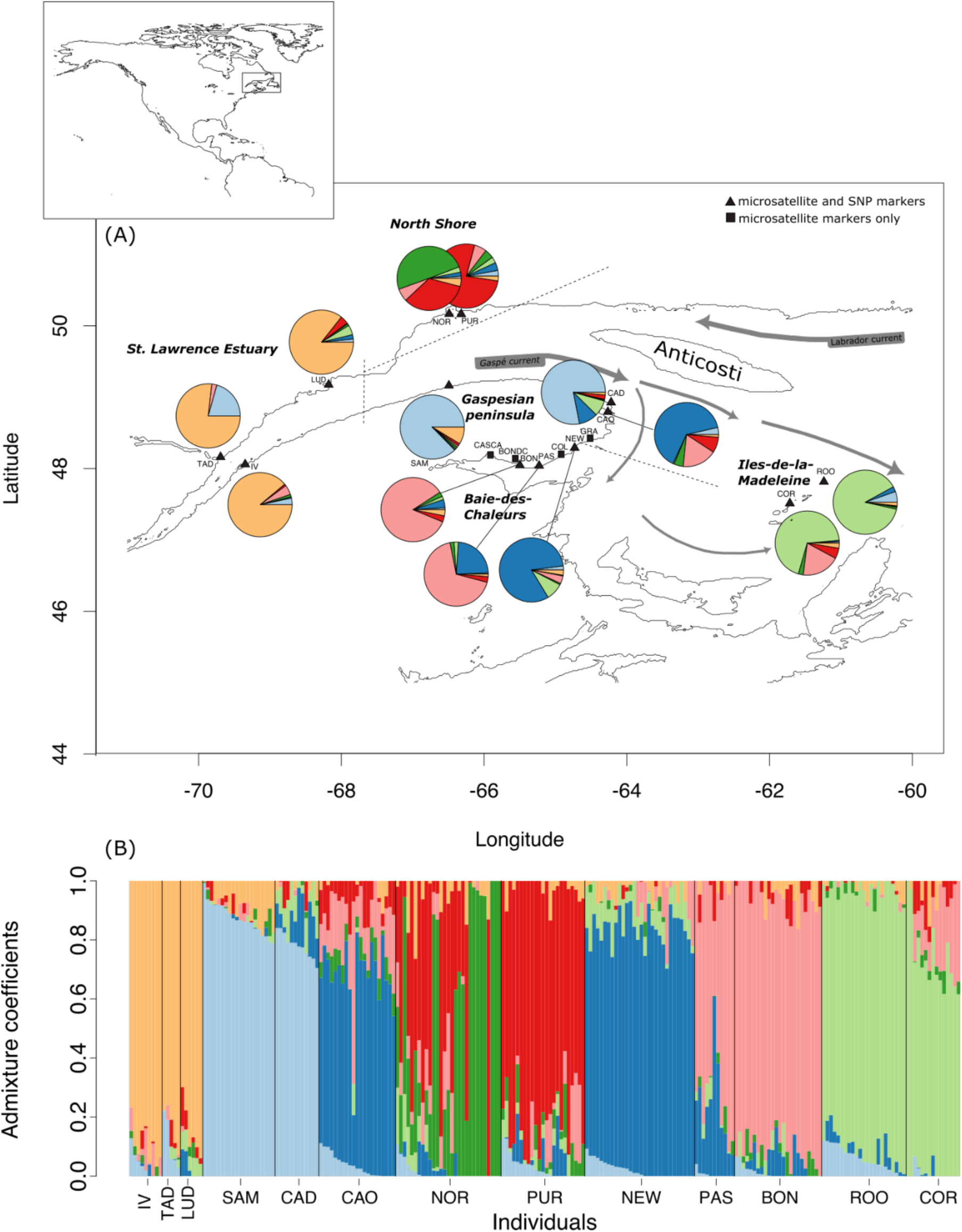
(A) Map of the sampling sites for *Saccharina latissima* in the Estuary and Gulf of St. Lawrence, Quebec, Canada. Pie charts correspond the relative proportion of the 7 putative ancestral populations inferred by Bayesian clustering of the genetic diversity at SNP markers. Individual clustering values are presented in (B).

## Materials and Methods

### Microsatellite markers

#### Sampling

Adult sporophytes of *S. latissima* (n = 480) were sampled throughout the Estuary and Gulf of St. Lawrence in 2018 and 2019. Thirty individuals were collected at each site in the Estuary: Tadoussac (TAD), Ile verte (IV), Pointe Ludger (LUD) and in the Gulf along the North Shore: Ile Grosse Boule (Purmer algaculture site : PUR), Sept-Iles (NOR), along the Gaspé Peninsula: Sainte-Anne-des-Monts (SAM), Cap Desrosiers (CAD), Cap-aux-Os (CAO), Grande-Rivière (GRA), in the Baie-des-Chaleurs: Newport (NEW), Colbourne (COL), Paspébiac (PAS), sporophytes from Bonaventure on growing lines (BONDC), Bonaventure (BON), and Cascapedia (CASCA) and around the Iles-de-la-Madeleine: Rochers-aux-Oiseaux (ROO) and Cormandière (COR) (Figure 1; Table 1). Two small pieces of tissues (1 cm^2^) were cut from the basal blades of individual sporophytes and placed in silica gels until DNA extractions. The site ‘PUR’ is located in the vicinity of the growing lines and ‘BONDC’ consist of sporophytes on the growing lines.

**Table 1.** Sampling site and their GPS coordinates of *Saccharina latissima* individuals in the St-Lawrence Estuary and Gulf.

| Site | Latitude | Longitude |
| --- | --- | --- |
| Tadoussac (TAD) | 48.133119 | -69.705842 |
| Ile verte (IV) | 48.034772 | -69.340821 |
| Pointe de Ludger (LUD) | 49.161609 | -68.173798 |
| Ste-Anne-des-Monts (SAM) | 49.130887 | -66.500031 |
| Pointe-Noire (NOR) | 50.163282 | -66.467896 |
| Ile Grosse Boule (PUR) | 50.151276 | -66.325630 |
| Cap-des-Rosiers (CAD) | 48.859972 | -64.190300 |
| Cap-aux-Os (CAO) | 48.840380 | -64.331696 |
| Grande-Rivière (GRA) | 48.392669 | -64.501934 |
| Newport (NEW) | 48.263541 | -64.735703 |
| Colborne (COL) | 48.181379 | -64.895210 |
| Paspébiac (PAS) | 48.017946 | -65.265587 |
| Bonaventure (BONDC) | 48.031149 | -65.479357 |
| Bonaventure (BON) | 48.041124 | -65.491133 |
| Casapédia (CASC) | 48.175107 | -65.918175 |
| Cormorandière (COR) | 47.496607 | -61.723938 |
| Rocher-aux-Oiseaux (ROO) | 47.806514 | -61.232300 |

#### DNA Extractions

Total genomic DNA was extracted from 5 to 10 mg of dry tissue using the Nucleospin 96 plant kit (Macherey-Nagel, Germany). The extractions were performed according to the manufacturer’s instructions, except that samples were left in the lysis buffer at room temperature for one hour rather than being heated to 65°C for 30 min. The extracted DNA was eluted in 200 μL of the supplied elution buffer and one additional wash with buffer PW1 and one additional wash with buffer PW2 to more effectively remove PCR inhibitors and polysaccharides.

#### Microsatellite loci amplification

Overall, 44 microsatellite loci for *S. latissima* were amplified. Twelve of these loci were previously described in Paulino *et al*. (2016) and 32 loci cloned from expressed sequence tag-derived (EST) by Guzinski *et al*. (2016). For the multiplexes with the 12 loci characterized by Paulino *et al*. (2016), PCR mixes were prepared following their instructions except for dNTPs (which were adjusted to 150 µM) and the Taq polymerase (adjusted to 0.5 U). For the multiplexes from the EST, amplifications were carried out in 20 μL reaction volumes with each reaction comprising 2 μL of DNA template diluted 1:50 (to help reduce PCR inhibitors), 2 mM of MgCl2, 1 X GoTaq® Flexi PCR buffer, 150 µM of dNTP, 400 nM of forward primer, 400nM of reverse primer and 0.5 U of GoTaq® Flexi (Promega Corporation, Madison, WI, USA). All PCR amplifications were performed using a T100TM Thermal Cycler (Bio-Rad Laboratories, Inc., Hercules, CA, USA) and parameters used are described in Paulino *et al*. (2016) and Guzinski *et al*. (2016) (for the EST multiplexes). Next, 2 μL of PCR product was added to 10 μL of loading buffer made up of 0.5 μL of the SM594 size standard (Mauger et al. 2012) and 9.5 μL of Hi-Di formamide, denatured at 95 °C for 3 min, and run in an ABI 3130 XL capillary sequencer (Applied Biosystems, USA). Genotypes were scored manually in Genemapper version 4.0 (Applied Biosystems) and verified by two readers.

#### Genomic markers

A total of 282 adult sporophytes of *Saccharina latissima* selected from 13 locations were sent to Diversity Arrays Technology (DArT) at the University of Canberra (Australia). Three sites were in the St. Lawrence Estuary (TAD, IV, LUD), two along the North Shore (PUR, NOR), three along the Gaspé Peninsula (SAM, CAD, CAO), three in the Baie-des-Chaleurs (NEW, PAS, BON), and two around the Iles-de-la-Madeleine (ROO, COR). Libraries were constructed according to Kilian et al. (2012). DArTseq is a proprietary genome complexity reduction-based sequencing technology that differs from other methods through its ability to select the predominantly active – low copy sequence – areas in a genome, which are the ones containing the most useful information. At the same time, DArTseq masks the lesser value, repetitive sequences. It does this through the application of a combination of restriction enzymes to fragment DNA samples in a highly reproducible manner. DArTseq complexity reduction method was used through digestion of genomic DNA with two restriction enzymes (*PstI* and *MseI*) and ligation of barcode adapters followed by PCR amplification of adapter-ligated fragments. Libraries were sequenced using single read sequencing runs for 77 bases by Hiseq2500. DArTseq markers scoring was achieved using DArTsoft14 which is an in-house marker scoring pipeline based on algorithms. Thirty replicates were produced to assess the reproducibility of the sequences and average reproducibility was 99%. Markers were aligned to the *Saccharina japonica* (SJ6l.1) genome.

### Data analysis

#### Microsatellite markers

The presence of sequencing errors like stuttering bands, allelic dropouts, and null alleles was verified with the MICRO-CHECKER software version 2.2.3 (Van Oosterhout et al. 2004). Monomorphic loci, loci with more than 30% of missing data, and loci with missing data for all individuals for a site were removed from the database. All individuals with more than three missing data per loci were also removed from the database. After selection, 22 loci were retained for 308 individuals. All locations were kept for analyses except CAO since it contained too many missing data. The number of individuals per site ranged from 9 to 28. Allelic richness was estimated in FSTAT, version 2.9.4 (Goudet 2005a).

#### Genomic markers

Initially, we received 68,971 single-row SNP markers from DArT Pty Ltd. These DArT loci were then filtered using the “dartR” library (Gruber et al. 2018) in Rstudio (version 4.2) (Rstudio 2015) to obtain only informative loci. Four filters were used. We removed (1) all the monomorphic loci (27,525 remaining SNP), (2) the loci with more than 0.5% of missing data (20,131 remaining SNP), (3) the duplicate loci (18,623 remaining SNP), (4) the individuals with more than 20% of missing data (removed 33 individuals). After recalculating loci call rate, a total of 6,582 SNP remained. The number of individuals ranged from 9 to 30 except for TAD that had only two individuals and was not considered for diversity indices estimations. Genetic diversity indices and population differentiation analyses were estimated only for sampling sites with a minimum sample size of 9 individuals.

#### Both marker types

Genetic diversity indices were estimated using the “hierfstat” package (Goudet 2005b). FIS were calculated together with these indices and the significance was calculated with 1000 iteration bootstraps. Genetic differentiation was estimated with the “hierfstat” package (Gruber et al., 2018) using fixation index FST (Weir and Cockerham 1984) and its statistical significance was tested by performing 1000 bootstraps.

A Principal Component Analysis (PCA) was performed to allow a visual interpretation of the genetic structure of the data using the DartR package version 2.9.7 (Gruber et al. 2018). The analyses of isolation by distance were tested using the in-water non-Euclidean paths as distance between sites. These data were retrieved using the *marmap* package version 1.0.10 (Pante and Bouhet 2013). Individual ancestry coefficients for both microsatellites and the SNP data were estimated using the sparse non-negative matrix factorization (sNMF) method (Frichot et al. 2014) as implemented in the R package LEA (Frichot & François, 2015). sNMF is robust against deviations from Hardy-Weinberg equilibrium and does not rely on any a priori population genetic assumptions (Frichot et al., 2014). This method offers comparable performance to Structure and ADMIXTURE while providing significantly faster run times for genomic datasets. The number of putative clusters (K) ranged between 1 and the number of sampling sites (depending on the marker) and for each K, 100 repetitions with 99 maximum iterations were carried out. Ten percent of the genotypes were masked at each run to compute the cross-entropy criterion. K with the smallest cross-entropy value and/or from which this parameter stabilized was selected. Analyses of molecular variance (AMOVA) were performed using the poppr R package (Kamvar et al 2015). Group discrimination was based on PCA results on SNP markers: Group1: IV, LUD, SAM, CAD; Group 2: NOR, PUR; Group3: NEW, BON, PAS, CAO and Group4: Iles-de-la-Madeleine (ROO and COR) and applied to both microsatellite and SNP data.

## Results

### Genetic diversity

Genetic diversity indices calculated with microsatellite markers were relatively low, with He values ranging from 0.249 to 0.339 and a mean number of alleles of 2.8 on average per site (Table 2, Figure 2a). Moreover, more than half of the sites had positive and significant FIS values. Only five sites were in Hardy-Weinberg equilibrium, the other ones showed heterozygosity deficiency. Genetic diversity indices using SNP indices were 10x lower, with expected heterozygosity (He) ranging from 0.0656 (IV) to 0.083 (PUR) (Table 2, Figure 2b). Populations from the Estuary and Gaspé Peninsula (CAD, SAM, IV, LUD) had lower heterozygosity than the rest of the sampling sites. Seven sites had positive FIS values, five of them also showed positive values at microsatellite markers. Sites from the Estuary had lower heterozygosity values than those from the Gulf at SNP but not at microsatellite markers.

**Figure 2.**
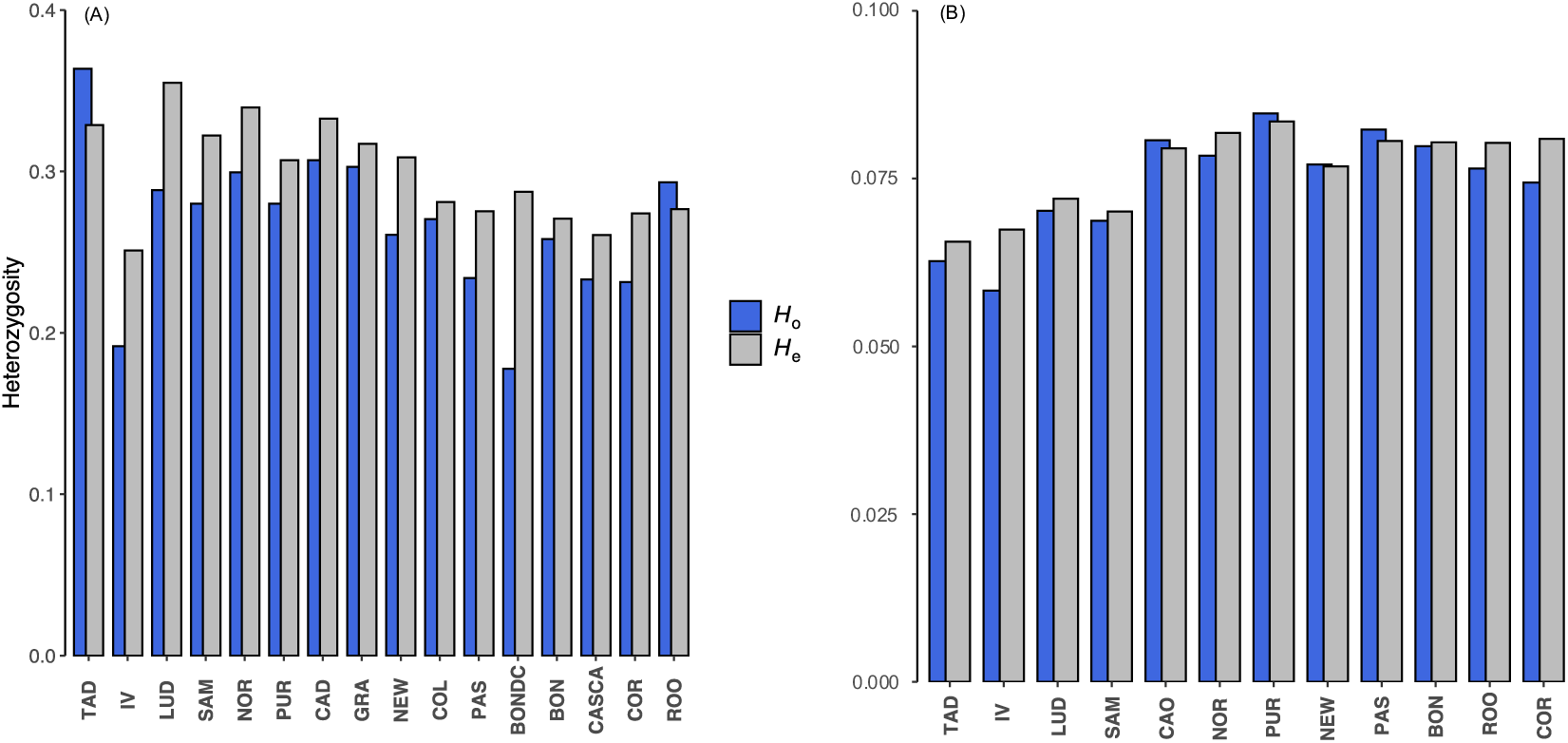
Heterozygosities (observed and expected) estimated from a) microsatellite markers and b) SNP and presented for each sampling site.

**Table 2.** Genetic diversity indices for microsatellite and SNP methods: Sample size (n), expected heterozygosity (He), mean allelic richness (AR), inbreeding coefficient values (FIS). Values in bold with an asterisk are significant.

| Microsatellites |  |  |  |  | SNPs |  |  |
| --- | --- | --- | --- | --- | --- | --- | --- |
| Site | n | He | AR | $F_{IS}$ | n | He | $F_{IS}$ |
| TAD | 14 | 0.330 | 1.68 | -0.106 | - | - | - |
| IV | 15 | 0.249 | 1.51 | **0.236\*** | 9 | 0.0656 | **0.0396\*** |
| LUD | 9 | 0.336 | 1.66 | 0.188 | 9 | 0.0671 | **0.1040\*** |
| SAM | 22 | 0.321 | 1.66 | **0.131\*** | 20 | 0.0719 | **0.0200\*** |
| NOR | 22 | 0.339 | 1.71 | **0.118\*** | 29 | 0.0890 | **0.0256\*** |
| PUR | 20 | 0.306 | 1.64 | **0.088\*** | 23 | 0.0833 | -0.0063 |
| CAD | 19 | 0.332 | 1.69 | **0.077\*** | 12 | 0.0701 | **0.0158\*** |
| CAO | - | - | - | - | 21 | 0.0795 | -0.0076 |
| GRA | 15 | 0.317 | 1.66 | 0.045 | - | - | - |
| NEW | 21 | 0.308 | 1.66 | **0.156\*** | 30 | 0.0764 | 0.005 |
| COL | 23 | 0.280 | 1.58 | 0.038 | - | - | - |
| PAS | 23 | 0.274 | 1.57 | **0.150\*** | 11 | 0.081 | -0.0141 |
| BONDC | 13 | 0.281 | 1.6 | **0.381\*** | - | - | - |
| BON | 23 | 0.270 | 1.58 | 0.047 | 24 | 0.0804 | 0.0102 |
| CASCA | 22 | 0.260 | 1.53 | **0.105\*** | - | - | - |
| COR | 21 | 0.273 | 1.56 | **0.155\*** | 13 | 0.0809 | **0.0611\*** |
| ROO | 26 | 0.277 | 1.58 | -0.060 | 25 | 0.0803 | **0.0368\*** |

### Genetic structure

The matrix of pairwise FST (Figure 3a) revealed low genetic differentiation among the 16 locations using microsatellite markers. Significant FST values were found between the Iles-de-la-Madeleine location (ROO) and most of the other sites and between the Sept-Iles site and many locations. *S. latissima* from Bonaventure (BON) used for algal culture showed small but significant genetic differentiation with individuals collected in NOR (located 1 km from the cultivation line on the North Shore). All pairwise FST obtained from SNP were significant with highest between the Iles-de-la-Madeleine and the Estuary (from 0.18 to 0.12) and lowest values among sites within the Baie-des-Chaleurs (from 0.01 to 0.08) (Figure 3b). Individuals from PUR (aquaculture site) showed little genetic differentiation from NOR individuals located 1 km from that site and were genetically differentiated from the Baie-des-Chaleurs site (BON) where meiospores were collected from 2014 to 2018. Mantel test revealed no significant isolation by distance at microsatellites (Figure 4a) but a significant relationship at SNP (Figure 4b). The PCA analyses from microsatellite markers revealed the presence of a single group composed of all 16 sites (Figure 5a). By contrast, PCA with SNP markers revealed significant structure, with Estuary (TAD, IV, LUD) and Gaspé Peninsula sites (SAM and CAD) forming the first group, North Shore sites (NOR and PUR) on the first axis (6.93% of total genetic variance) forming a second group. The Baie-des-Chaleurs sites (BON, PAS, NEW), CAO, and the Iles-de-la-Madeleine sites (ROO and COR) separated along the second principal component (4.71%) (Figure 5b) formed the third group.

**Figure 3.**
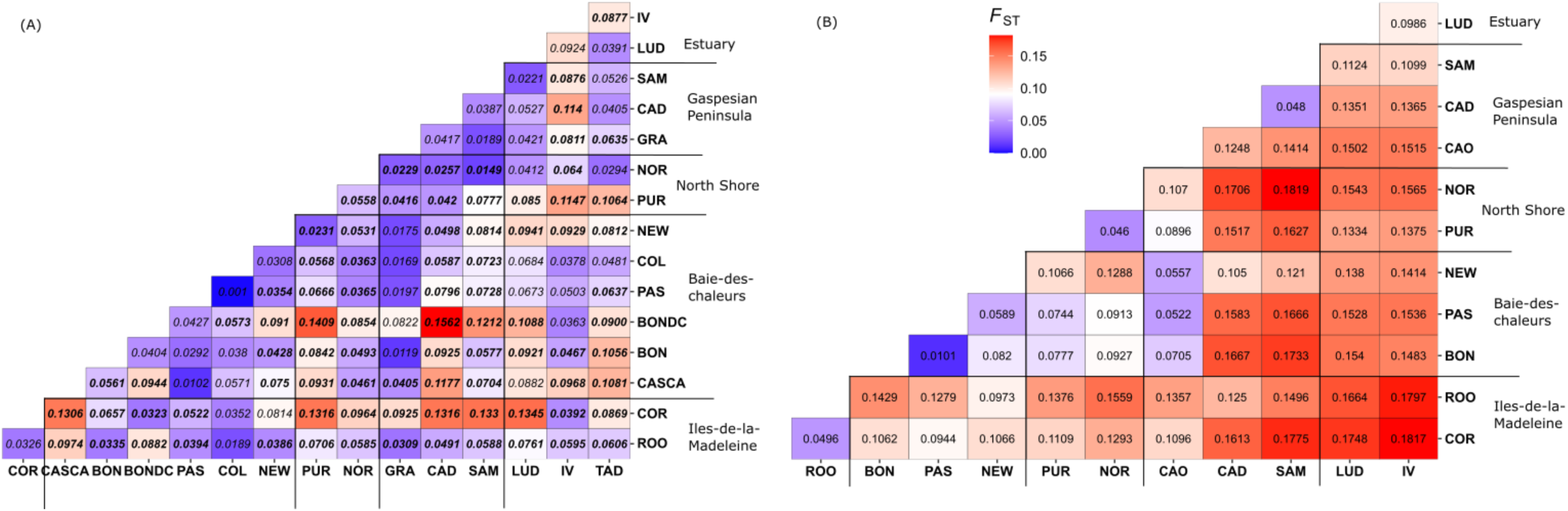
Matrix of pairwise FST estimated from a) microsatellite and b) SNP markers.

**Figure 4.**
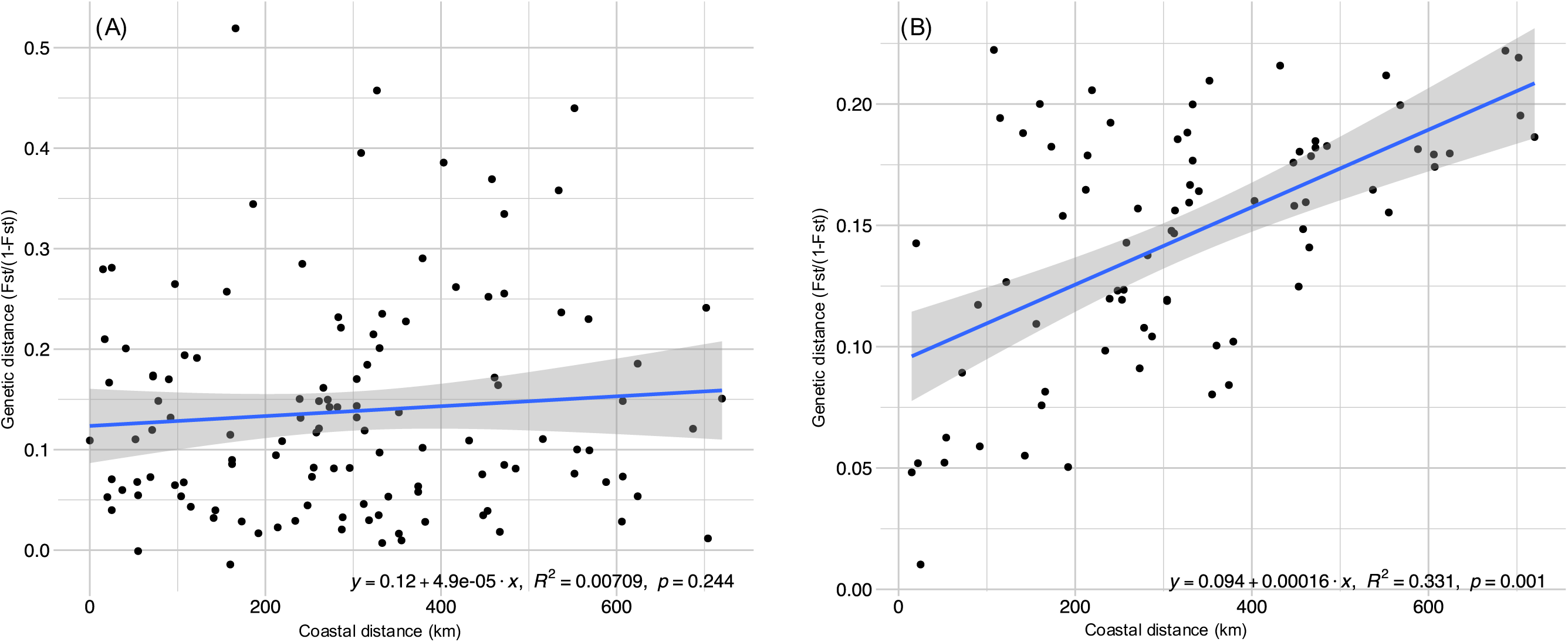
Mantel test plot performed using a) microsatellites, and b) SNP markers displaying the correlation between genetic and coastal distances among sugar kelp populations. Correlation coefficient r and p-value are shown.

**Figure 5.**
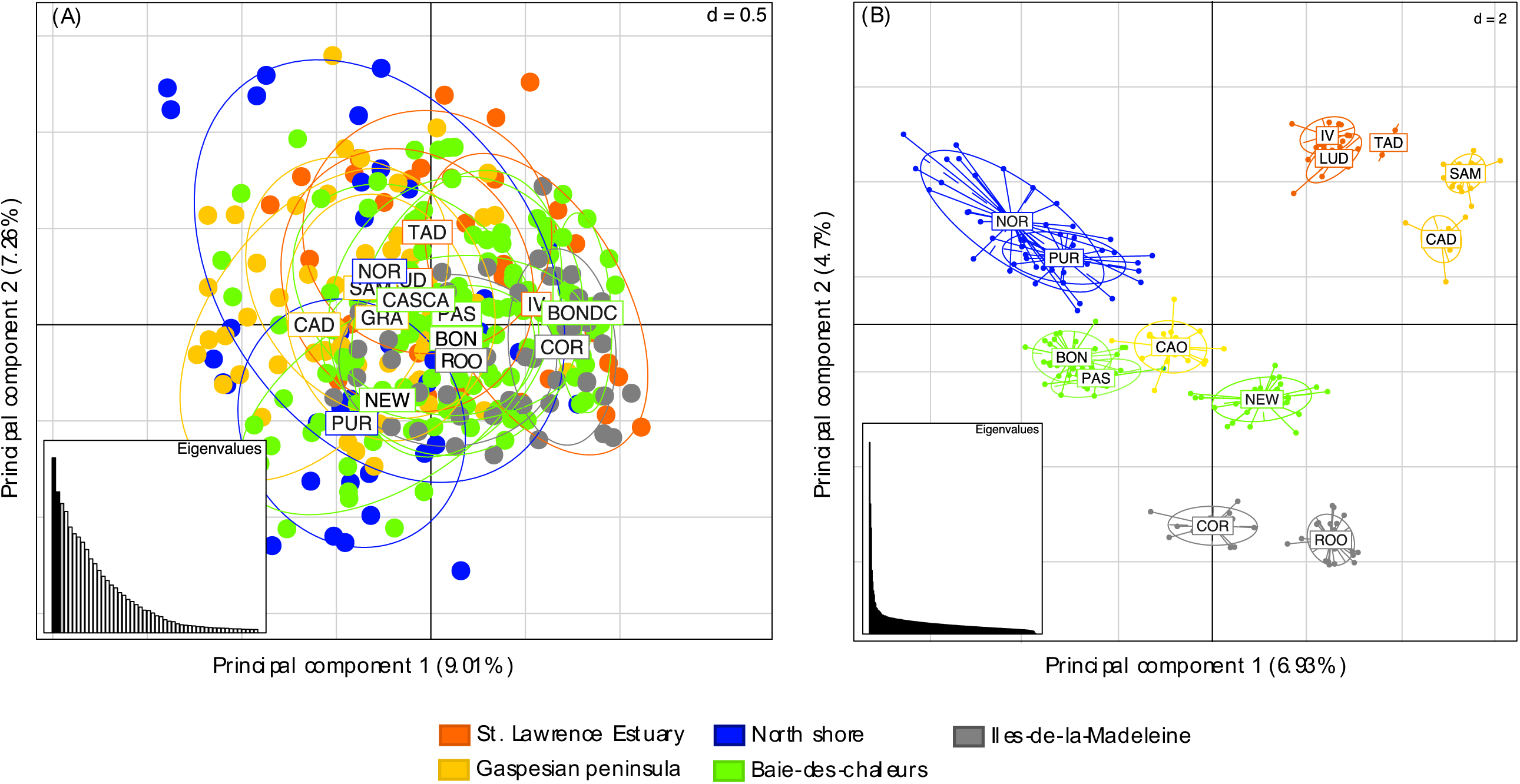
Principal Component Analysis (PCA) results based on a) microsatellite, and b) SNP markers from individuals of *Saccharina latissima* (n=228) collected in the Estuary and Gulf of St. Lawrence. Each point represents an individual, and each color corresponds to one site.

Population structuration analyses determined that one ancestral population was the most likely scenario with microsatellite markers, with the minimum cross-entropy recorded at K=1. When using the SNP markers, the most probable number of ancestral populations was 7, after which the cross-entropy criterion reached a plateau (Figure 1). The estuarian sites (IV, TAD and LUD) all shared a common major ancestral cluster, the Gaspesian sites (SAM and CAD) shared another one, the sites from Iles-de-la-Madeleine (ROO and COR) formed a third cluster (Figure 1). The CAO site, separated from CAD on the Gaspé peninsula by a narrow land formed a group with NEW. The sites located deeper in the Baie-des-Chaleurs (PAS and BON) formed another cluster. The sites from the north shore (NOR and PUR) were separated into distinct clusters.

AMOVA analyses explained only 0.71% of the variation at microsatellite loci (this percentage was not significant) whereas the variance explained by the 4 groups at SNP markers accounted for more than 7 % of the total variation and was significant (p=0.01) (Table 3).

**Table 3.**
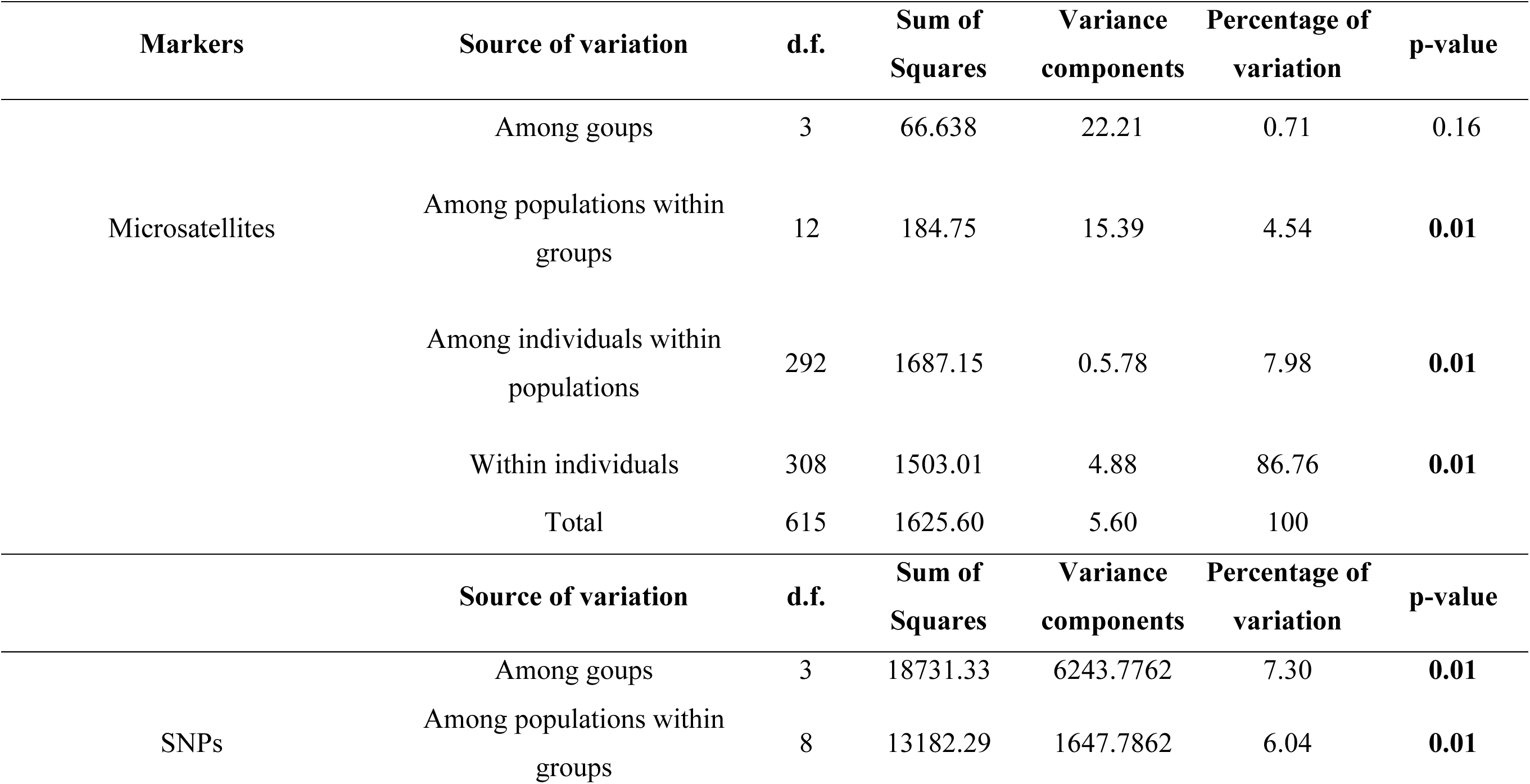
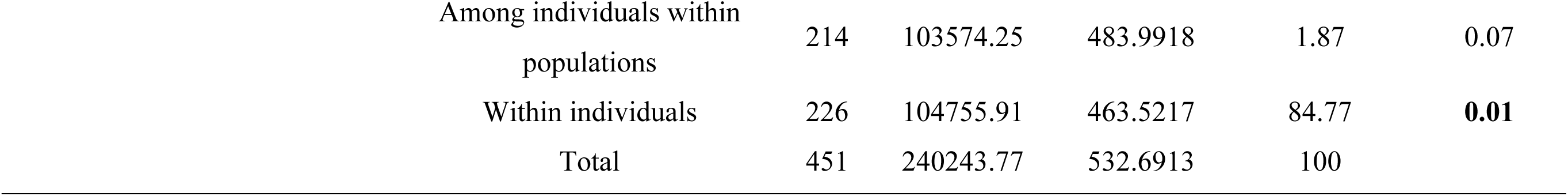
Analysis of molecular variance (AMOVA) using four groups based on group discrimination revealed by PCA on SNP markers: roup1: IV, LUD, SAM, CAD; Group 2: NOR, PUR; Group3: NEW, BON, PAS, CAO and Group4: Iles-de-la-madeleine (ROO and COR).

## Discussion

This study is the first to examine patterns of genetic diversity and structure in wild populations of *Saccharina latissima* from the St. Lawrence Estuary and Gulf. Results showed low genetic diversity in *S. latissima* from the St. Lawrence seaway with half of the sites showing high FIS indices and a low number of effective alleles per locus. Observed low levels of genetic diversity may be due to bottlenecks following postglacial colonization in the St. Lawrence River. A similar study in Maine, USA found higher levels of allelic richness (3.6 and 4.2) with only six microsatellite markers and no inbreeding (Breton et al. 2018). A worldwide survey using 12 microsatellite loci reports that the Eastern Atlantic (Europe) populations are more genetically diversified than the Western Atlantic (US and Canadian Coasts) and the Pacific populations (Neiva et al. 2018). Heterozygosity levels were lower in NW Atlantic *S. latissima* populations and several sites had high inbreeding levels (Neiva et al. 2018). Three of our sites were sampled in their study (NOR, NEW, and BON) and had an average allelic richness of 3.9, that was similar to other populations from the NW Atlantic but higher than our current values (around 3). In contrast to our present results comparing the same sites, Neiva et al. (2018) did not detect any inbreeding at our three locations but at several other NW sites including the Canadian Maritimes. Allelic richness was twice as high in NE Atlantic populations compared to our results for the same set of microsatellite loci (Guzinski et al. 2016). Higher levels of genetic diversity in the NE than in the NW populations were also found in *Laminaria digitata* (Neiva et al., 2020). Guzinski et al. (2020) found higher levels of genetic diversity at SNP (from ddradseq) in the northernmost localities of Norway and Scotland (Ho of 0.05) than in more southern populations located in Portugal (Ho of 0.030). Among studies comparisons of genetic diversity indices are more difficult with SNP than with microsatellites since the type of sequencing can differ, the sequencing coverage as well as the various filtrations on the datasets. No studies have directly compared diversity indices at SNP for *S. latissima* populations from NE and NW Atlantic.

Recurrent bottlenecks following Pleistocene glaciation may have affected the genetic diversity of sugar kelp in the St. Lawrence. During this period, 25,000-18,000 years ago, the ice sheet covered North America, from the Arctic to approximately New York, and the St-Lawrence was wholly sealed in ice (Maggs et al., 2008). After this period, many species, such as *S. latissima*, recolonized ice-free areas. It has been assumed that populations of the NW Atlantic were composed of individuals that migrated across the Atlantic Ocean from Europe following the last glaciations (Lee, 1973; Taylor, 1957, Luttikhuizen et al. 2018). Neiva et al. (2018) suggested that NW and NE lineages evolved in the Atlantic and split into two groups prior to the late glacial maximum. The consequences of glaciations were harsher in the NW Atlantic due to the paucity of rocky shores resulting in lower genetic diversity in members of this phylogroup. Few studies have examined recolonization patterns in species from the St. Lawrence. The anadromous rainbow smelt (*Osmerus mordax*) recolonized the St. Lawrence Estuary from two refugia (Bernatchez, 1997). The Mysid *Neomysis americana* includes two different lineages one that would have colonized the St. Lawrence via a southern refuge around Georges Bank with a migration route via the Champlain Sea and another one with an origin east of the Grand Banks, Newfoundland (Cortial et al., 2019) Few studies have compared genetic diversity among the Estuary and the Gulf of St. Lawrence in species that do not actively disperse. Our results showed low levels of genetic diversity, suggesting recolonization from a small number of individuals and a single refugium. Additional studies examining mitochondrial genomes would be needed for confirmation. Environmental factors may also affect genetic diversity levels. Populations from the more brackish environment (lower Estuary) were less genetically diverse than those from the Gulf at genomic markers. A similar result was found for *Zostera marina* populations from the same area and sequenced with the same technique (Dartseq) (Treillefort 2023). Nielsen et al. (2016) found that brackish populations of *S. latissima* in Denmark were less genetically diverse than marine populations and likely more vulnerable to climate change due to their genetic isolation.

Very weak genetic structure was revealed for the 22 microsatellite markers in comparison with SNP. The use of genomic markers allowed an increased resolution and the detection of restriction to gene flow at major barriers in the St. Lawrence seaway. Spatial analysis of genetic distance using SNP supported the influence of isolation by distance (IBD) on population differentiation across the whole region. The dataset also supported the presence of important hierarchical structuring. The Gaspé current flows along the south shore of the estuary and along the Gaspé peninsula thus explaining the first and second cluster comprising individuals from the lower estuary (IV, TAD, LUD) and the Gaspé peninsula (SAM, CAD). The isolation of North Shore populations (NOR and the algal culture site, PUR) can be explained by the presence of a gyre near Anticosti Island (Archambault et al. 2017). These two sites were found to be distinct from one another despite the low FST values between them. Additional samples at sites located to the east of this location would help to understand the genetic divergence between these sites. The populations (BON and PAS) which are located to the west of the Baie-des-Chaleurs clustered together whereas NEW and CAO occupying the eastern part of that area formed a distinct cluster. The Gaspé current flowing from the Gaspé Peninsula to the Iles-de-la-Madeleine (Savenkoff et al, 1996) likely prevents mixing between these sites and the sites from the Gaspé peninsula. The Iles-de-la-Madeleine (ROO, COR) are further isolated due to distances from the rest of the sites. In Maine, *S. latissima* shows genetic differentiation within a few kilometers, four groups were detected with microsatellites along only 210 km of shoreline (Breton et al. 2018). In eastern USA, Gulf of Maine and southern New England populations were separated by Cape Cod, a peninsula that appears to be a biogeographic barrier to the gene flow in this area (Mao et al. 2020). In comparison, the Gaspesian peninsula could be a barrier to the gene flow separating the Estuary, the North Shore, and the Gaspesian coast from the Baie-des-Chaleurs.

Macroalgae farming has been in operation at very small scale since 2010 in the St. Lawrence. Our study revealed that wild population of the Baie-des-Chaleurs (BON) where the broodstock is collected is genetically distinct from wild population of Sept-Iles. The North Shore population where the aquaculture has recently been in operation (PUR) does not differ genetically from the wild population (NOR) located 1 km away. This suggests an absence of genetic contamination from the seed broodstock. As sea farming has not been practiced intensively, our results are not surprising. The use of locally sourced seeding material has been advocated to reduce the risk of genetic pollution and depression from introduced non-local strains (Yarish et al., 2017). Low levels of introgression have been reported in naturalised populations of *Undaria pinnatifida* from nearby farm sites in France (Grulois et al. 2011). A recent study comparing 24 wild and three farmed *S. latissima* populations at 21 microsatellite loci revealed no significant genetic differentiation between them, reflecting the current cultivation practices of using local meiospores in Brittany, France (Jaugeon et al. submitted). In China, little signs of gene flow from farm to wild has been reported in U. *pinnatifida* (Li et al., 2020). As genetic diversity appears limited in our populations and genetic structuration is high, it would be important to take this into consideration for selection programs and if farming activities intensify. The introduction of genetic diversity through the crossing of divergent populations of *S. latissima* has been demonstrated to result in heterosis, leading to increased yield (Cohen et al. 2025). The recent report of sporeless sprorophyte production could alleviate the problem of genetic contamination and enable the production of desirable traits in cultivated *S. latissima* (Vissers et al. 2024). However, such a strategy raises the question of farmers’dependance on seed producers. To promote more independence on seed producers, it is important to maintain/explore alternative options such as local strains adapted to their specific populations.

This study provides the first population genomic survey of an important kelp foundation species in the gulf of St.Lawrence. This portrait is greatly needed to understand the effects of global changes and transfer of genetic material among the main groups of this key species in the St. Lawrence. The low diversity observed may infer that populations could be less resilient to climate change or farming intensification. Future studies should focus on identifying genomic regions associated with temperature and salinity tolerance. This would help to understand the evolutionary potential of this species facing future disturbances as well as to create aquaculture strains with sufficient tolerance to future perturbations. The important genetic structure should serve as a guideline for the management, conservation, and cultivation of sugar kelps.

## Acknowledgements

This study was supported by an aquaculture Collaborative Research and Development Program (ACRDP) to Tamara Provencher, Rénald Belley, and France Dufresne and core founding from the CNRS and Sorbonne Université to the IRL EBEA 3614. Marie Treillefort and Sabrina Le Cam received internships with Merinov funded by Mitacs acceleration grants. We acknowledge travelling grants from Institut-France-Québec for a stay in the laboratory of Myriam Valero in Roscoff. We thank the various people and industrial partners who helped with sampling sugar kelps. We are also grateful to the Biogenouest genomics core facility (Genomer Plateforme génomique) at the Station Biologique de Roscoff for their technical support.

